# BioDH: A reversible data hiding framework for secure hiding sensitive data in biological data

**DOI:** 10.1101/2025.10.28.685003

**Authors:** Zhang Yu, Lin Yijie, Dong Haozhe, Fu teng

## Abstract

With the rapid advancement of single-cell RNA sequencing (scRNA-seq) technologies, the open sharing of massive high-dimensional biological data has become a key driver in advancing precision medicine and fundamental research. However, while promoting data accessibility, ensuring effective protection of sensitive data has emerged as a critical challenge in bioinformatics. To address the issue, we propose BioDH (Biological Data Hiding) framework, a reversible information hiding framework tailored for biological data, aiming to simultaneously ensure data security and preserve scientific utility. The framework employs optimal lossless compression and strictly controls numerical perturbations to maintain precision. Upon extraction, both the secret information and the original data are perfectly recovered, achieving true reversibility. Validation on real-world datasets shows exceptional fidelity: when embedding capacity is 1.25 bit per byte, maximum and mean absolute errors are 8.54E-04 and 8.90E-11, respectively; PCA reveals a correlation of 1.0; UMAP testing exhibits no structural distortion; Mantel tests and clustering analyses (ARI = 1.0) confirm preservation of high-dimensional topology and cell subpopulations. All metrics surpass biological compatibility thresholds, indicating no detectable impact on downstream analyses.

## 1 Introduction

With the rapid advancement of high-throughput sequencing technologies, particularly single-cell RNA sequencing [1], biomedical research has entered an unprecedented big data era. The sequencing data not only possesses extremely high scientific research value but also contains abundant individual sensitive data, such as patients’ genetic backgrounds, disease statuses, and even personal identifiers. Under China’s Personal Information Protection Law and Data Security Law, biometric data is explicitly listed as sensitive personal information. Their collection and sharing require the highest-level security safeguards; unauthorized processing can trigger heavy administrative fines, civil damages and up to years’ imprisonment [2–4].

How to effectively prevent the leakage of sensitive data while promoting scientific collaboration and data openness has become a key issue urgently needing resolution in the current field of bioinformatics. Traditional data protection methods face critical limitations. To bridge this gap, this paper proposes BioDH(Biological Data Hiding), the first reversible data hiding framework for scRNA-seq data. BioDH ensures invisibility, reversibility, zero size expansion, and seamless workflow integration and enables secure, auditable, and analysis-preserving data sharing without signaling data sensitivity.

Compared to existed approaches, our work makes the following core contributions. (1) The first framework for reversible data hiding in scRNA-seq data is presented. By directly embedding sensitive information into gene expression values, covert transmission is achieved while enabling perfect, lossless recovery of the original data; (2) A dedicated DhStruct data structure is designed to integrate optimally compressed payloads, metadata, and control fields, ensuring both reversibility and high efficiency in data processing; (3) The biological utility of the data containing hidden information is preserved. Using the EMPR(Embedding with Modulo oPeration Replace) embedding algorithm, the stego data maintains statistical and biological consistency with the original data; (4) The complete source code and supplementary materials [5] are publicly released to support open, collaborative, and reproducible research in bioinformatics.

The paper is organized as follows. Section 1 introduces the research background, core challenges, motivations, and key contributions. Section 2 provides a systematic review of advances in privacy protection for bioinformatics, biological data steganography, and compression-steganography integration. Section 3 presents the BioDH framework, detailing data preprocessing, dynamic compression coding, EMPR reversible embedding, information extraction, and data restoration mechanisms, along with strategies for biological compatibility validation. Section 4 evaluates the framework on public datasets through comparative experiments. Section 5 summarizes the work.

## 2 Related Work

Traditional Data Protection Methods for securing bioinformatics data primarily focus on access control, data anonymization, and encryption-based storage or transmission.1) Access control [6, 7];2) Data anonymization [8, 9];3) Encrypted storage and transmission such as AES [10] and RSA [11]. These methods ensure confidentiality by transforming data into an unreadable format without the correct decryption key. Despite their strong security guarantees, encrypted data cannot be directly analyzed—requiring decryption before any computational task, which undermines the biological utility of the data. Moreover, the presence of encryption itself signals data sensitivity, potentially attracting adversarial scrutiny or targeted attacks.

Contemporary privacy-preserving technologies fall into the following categories: 1) Homomorphic encryption [12, 13]: enables computation on encrypted data, protecting privacy during processing. However, it incurs high computational overhead and offers limited support for complex analyses; 2) Differential privacy [14–16]: protects individual privacy by adding calibrated noise to data or query results, preventing inference about any single subject. It introduces inevitable accuracy degradation, potentially compromising downstream analyses that require high precision; 3) Federated learning [13, 15, 17]: enables collaborative model training across multiple parties without sharing raw data, which remain local. It suffers from high communication costs, slow convergence, and substantial computational requirements for participants.

Data hiding technology [18–20] offers a new perspective for protecting biological data, with main applications including: 1) Medical image steganography: embedding patient information, copyright tags, or diagnostic reports into medical images. These studies mostly adapt conventional image steganography techniques, but the pixel-matrix carrier fundamentally differs from gene expression matrices in structure and semantics; 2) DNA sequence steganography operate at the molecular level but typically require synthesizing or altering actual DNA sequences, entailing high costs and ethical concerns; 3) Genomic data watermarking approaches primarily target discrete genotype data rather than continuous gene expression values, and often lack reversibility.

Recently, with the development of modern compression algorithms such as zlib, gzip and zstd [21], their compression applications in security contexts have gained increasing attention. In bioinformatics, data compression is primarily employed to reduce storage space and transmission bandwidth. However, research specifically leveraging compression to enhance data hiding efficiency remains scarce, particularly lacking systematic performance comparisons of various modern compression algorithms in the context of biological data steganography.

In summary, significant gaps remain in current research: a lack of specialized data hiding methods tailored for scRNA-seq data; an overreliance on irreversible watermarking, with few frameworks ensuring reversibility without compromising scientific utility; insufficient systematic evaluation of the impact of embedding on downstream biomedical analyses; and a scarcity of open-source implementations, limiting scalability and reproducibility. The proposed BioDH framework is designed to fill these gaps.

## 3 Methodology

### 3.1 BioDH framework

The proposed BioDH reversible data hiding framework designed for scRNA-seq data is shown in Fig. 1. BioDH supports the covert integration of sensitive information into gene expression matrices without compromising their scientific utility. The BioDH framework comprises the following modules: data preprocessing, compression encoding, embedding, file writing, information extraction, data restoration, and compatibility verification.

**Fig. 1.**
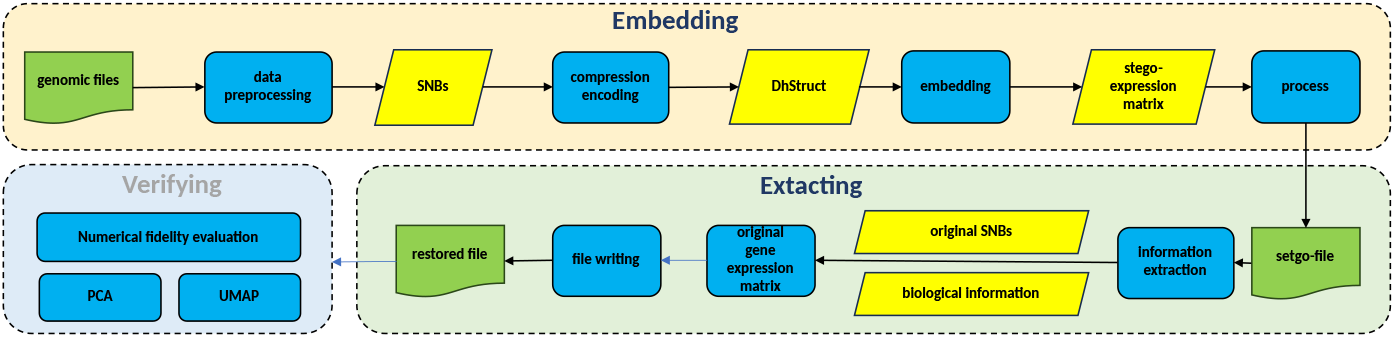
BioDH framework

The data preprocessing module reads genomic files and extracts the nonzero floating sequences from the matrix. These values are streamed for encoding, forming the basis of the data hiding process. The compression encoding module first extracts a specified number of bits (SNB) from the least significant digit of the mantissa of a floating number. Then, an optimal lossless compression algorithm is applied to efficiently compress the original SNB bitstream. The compressed data is organized into a specially designed DhStruct format. The embedding module then performs sequentially embedding the bits of the constructed DhStruct bitstream into the original data, The file writing module writes the modified data into a stego-file.

By reverse-parsing the DhStruct format, the information extraction module accurately retrieves the embedded information and the original SNB sequence. Using the SNB sequence recovered by the extraction module, the data restoration module reconstructs the original gene data. To evaluate consistency before and after embedding, the compatibility verification module performs comprehensive assessments using multiple metrics.

### 3.2 Data embedding process

The data embedding process and algorithm (EMPR)is shown in Algorithm 1. First, the input File is read, and the gene expression matrix is extracted. Nonzero or biologically meaningful gene expression values from the matrix are selected to form a significant floating list, denoted as FList. Next, each floating number in FList is traversed, and the SNB of the mantissa in its binary representation are extracted, forming the original bit sequence SNBList.

SNBList is compressed using a lossless compression algorithm. An optimal algorithm (e.g., zlib, Bz2, gzip, Zstd etc.) is selected based on evaluation of compression ratio. The compressed bitstream is denoted as SNBListComp. The choice of algorithm is recorded as a single-byte identifier within the data structure, ensuring correct decompression during extraction. This compressed bitstream, along with auxiliary metadata, is then structured into the DhStruct format for subsequent embedding. The DhStruct structure contains the following fields:(1) biological information length (32 bits), indicating the byte length of the original biological data;(2) algorithm identifier (8 bits), specifying the compression method used;(3) compressed data length (32 bits);(4) auxiliary information length (32 bits);(5) biological information content (variable-length);(6) compressed SNB data (SNBListComp);(7) optional auxiliary information. The DbStruct is serialized into a bitstream (DbStructBS).

Next, the system traverses the floating numbers in FList again, replacing the SNB of the mantissa in each number’s binary representation with modulo operation on DhStructBs, following a strategy of modular calculation on the lowest specified bits. Finally, the modified gene expression matrix is written back into a stego-file, completing the embedding process.

#### Algorithm 1

EmbedSecretWithEmpr

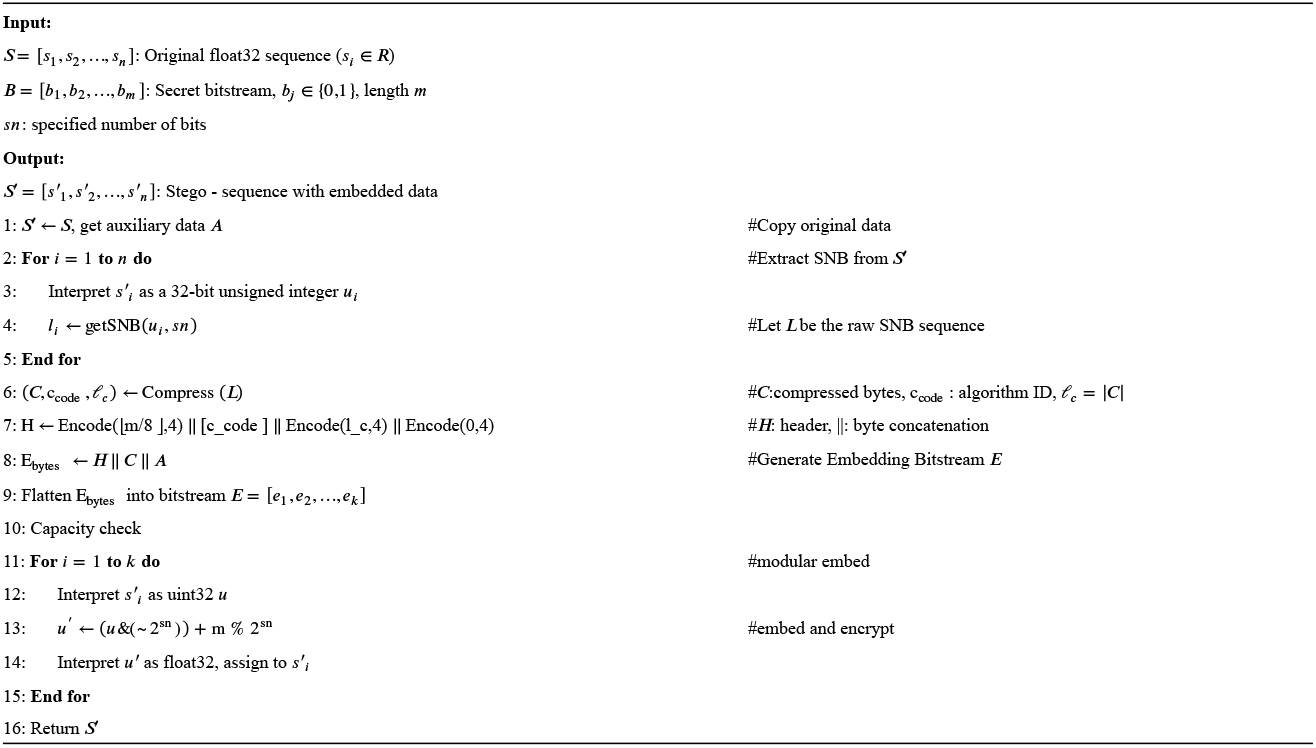

### 3.3 Data extraction process

The data extraction process recovers the original embedded biological information from the stego-file and enables full reconstruction of the original gene expression matrix, preserving its scientific utility. The complete extraction workflow is shown in Algorithm 2.

First, the stego-file is read and its gene expression matrix is loaded. Since the embedding operation only modifies the non-zero floating values, the modified floating sequence FListMod is extracted directly from the data. This sequence has the same length as the original FList, but the SNB of some values have been replaced with the hidden data. Next, each floating number in FListMod is parsed into 32-bit binary representation, and the SNB of the mantissa is extracted. These bits are concatenated in order to form a continuous bitstream which is the encoded form of the embedded DhStructBs. Subsequently, the DhStruct is parsed from the bitstream according to the predefined format. It sequentially extracts (1) the biological information content,(2) the compressed SNB data block (SNBListComp), and (3) the auxiliary information.

The compressed SNBListComp is then decompressed using the algorithm specified by the identifier, reconstructing the original SNB bitstream. This recovered SNB sequence is used to restore the original SNB sequence with modulo operation. The restored values are then placed back into the correct positions, fully recovering the original gene data and thus the restored file.

#### Algorithm 2

ExtractSecretWithEmpr

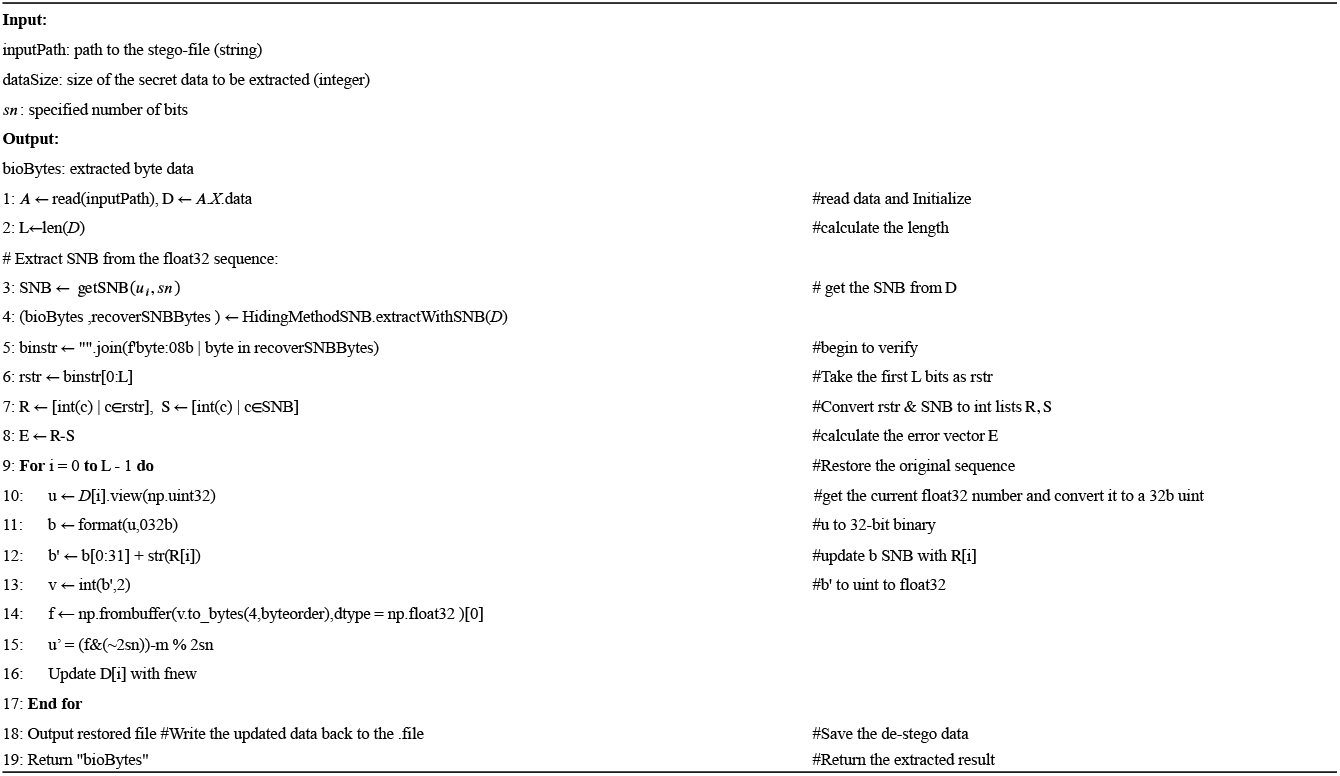

### 3.4 Biological compatibility validation process

To comprehensively evaluate the biological impact of information embedding on scRNA-seq data, this study establishes a multi-dimensional compatibility validation framework covering three aspects: numerical fidelity, clustering consistency evaluation, and nonlinear dimensionality and visualization consistency.

#### Numerical fidelity evaluation

To verify the correctness of the embedding and extraction processes, this paper extracts the embedded secret information and compares it with the original information. Meanwhile, the dataset is reconstructed from the stego-file, and the reconstructed dataset is compared with the original dataset. Additionally, the original dataset and the dataset with embedded information are compared to compute the maximum relative error and the mean error. Let the original data matrix be *A* = [*a*_*ij*_] and the stego-matrix be *B* = [*b*_*ij*_], where i=1,…,m and j=1,…,n, and m×n is the total number of elements in the matrix. The maximal and average absolute difference are computed as *diff*_*max*_ = max_*i,j*_(|*a*_*ij*_ − *b*_*ij*_|) and 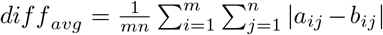 If *diff* is below a predefined threshold (*ε* = 0.001), the numerical changes are considered negligible and within the acceptable precision range for biological data.

#### Clustering consistency evaluation

Cell subpopulation identification is a core task in biological data analysis, directly Cell subpopulation identification is a core task in biological data analysis, directly influencing downstream biological interpretation. To assess whether information embedding disrupts cellular population structure, this study conducts a systematic validation at both the dimensionality reduction and clustering levels. At the dimensionality reduction stage, principal component analysis (PCA) [22] is performed, retaining the top principal components that capture the major expression patterns. The Pearson correlation coefficient between corresponding principal components of the original and embedded datasets is computed to evaluate the fidelity of global expression structure. The Adjusted Rand Index (ARI) [23]between clustering labels of the original and stego data is calculated to quantitatively assess the consistency of cell subpopulation assignments. A threshold (e.g., (*S*_*a*_*ri >* 0.98) is set to define high consistency between pre- and post-embedding clustering results.

#### Nonlinear dimensionality reduction and visualization consistency

To further validate the fidelity of high-dimensional manifold structures before and after information embedding, UMAP (Uniform Manifold Approximation and Projection) [24] is employed as the nonlinear dimensionality reduction method to assess consistency between the original and embedded datasets in the lowdimensional embedding space. UMAP constructs a fuzzy neighborhood graph in the high-dimensional space and optimizes its topological match in the lowdimensional space.

## 4 Experiments and Results

### 4.1 Experimental setup

This study is conducted on realscRNA-seq datasets, using publicly available data from the Human Cell Atlas (HCA) [25] as the primary input. For the experiments, some files in Annex 3(available at [5]) are used to test the embedding and extraction performance, ensuring the method’s applicability in realistic and complex biological contexts. The number of bits in SNB is set to 1, 2, 3, 4, 5. Two core functions were designed and implemented: ‘embedDataInH5ad()’ and ‘extractDataFromH5ad()’ using the algorithms described above. The former embeds secret message into the floating values of the gene expression matrix. The latter follows the inverse algorithmic path using the same DhStruct structural to precisely extract the hidden message and reconstruct the original expression matrix without requiring external keys.

### 4.2 Embedding capability

In the current experiment, up to 5 bits can be embedded in a single floating-point number. Since a floating-point number occupies 4 bytes, the embedding capacity can reach 1.25 bits per byte.

### 4.3 Numerical fidelity evaluation result

The experimental results are partly summarized in Table 1(more results are available at Annex2 [5]). The extracted secret information was completely identical to the original information, indicating that the information can be successfully embedded and accurately extracted. The reconstructed dataset was also completely consistent with the original dataset, demonstrating that the proposed data hiding method is fully reversible.

The experimental results show the maximum and mean absolute difference of 8.54E-04 and 8.90E-11, both of which are significantly below the predefined tolerance threshold. This indicates that the numerical perturbation introduced by the embedding process is extremely small and does not induce any substantive deviation. Therefore, the gene data before and after embedding are deemed to maintain high numerical fidelity, preserving the integrity required for downstream quantitative analyses.

### 4.4 Clustering consistency results

PCA was performed on both the original and stego datasets with the first 2 principal components retained. The result showed a PCA correlation coefficient of around 1.0, indicating that the embedding operation has no discernible impact on the overall covariance structure of gene expression. The result data show in Fig. 2.

**Fig. 2.**
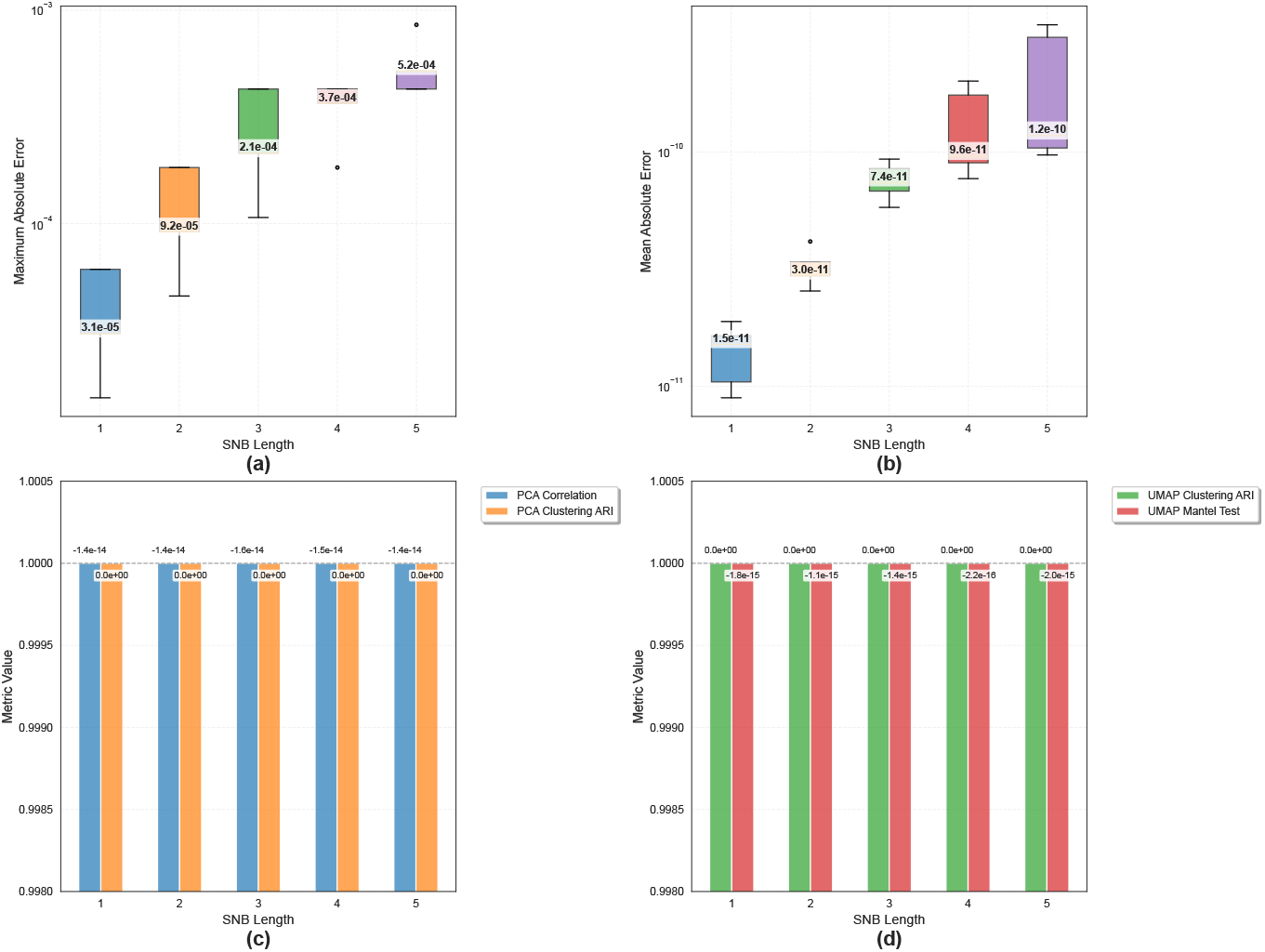
BioDH Framework Performance Analysis (a) Maximum Absolute Error vs SNB Length (b) Mean Absolute Error vs SNB Length (c) PCA-Based Biological Consistency Metrics vs SNB Length (d) UMAP-Based Biological Consistency Metrics vs SNB Length

**Fig. 3.**
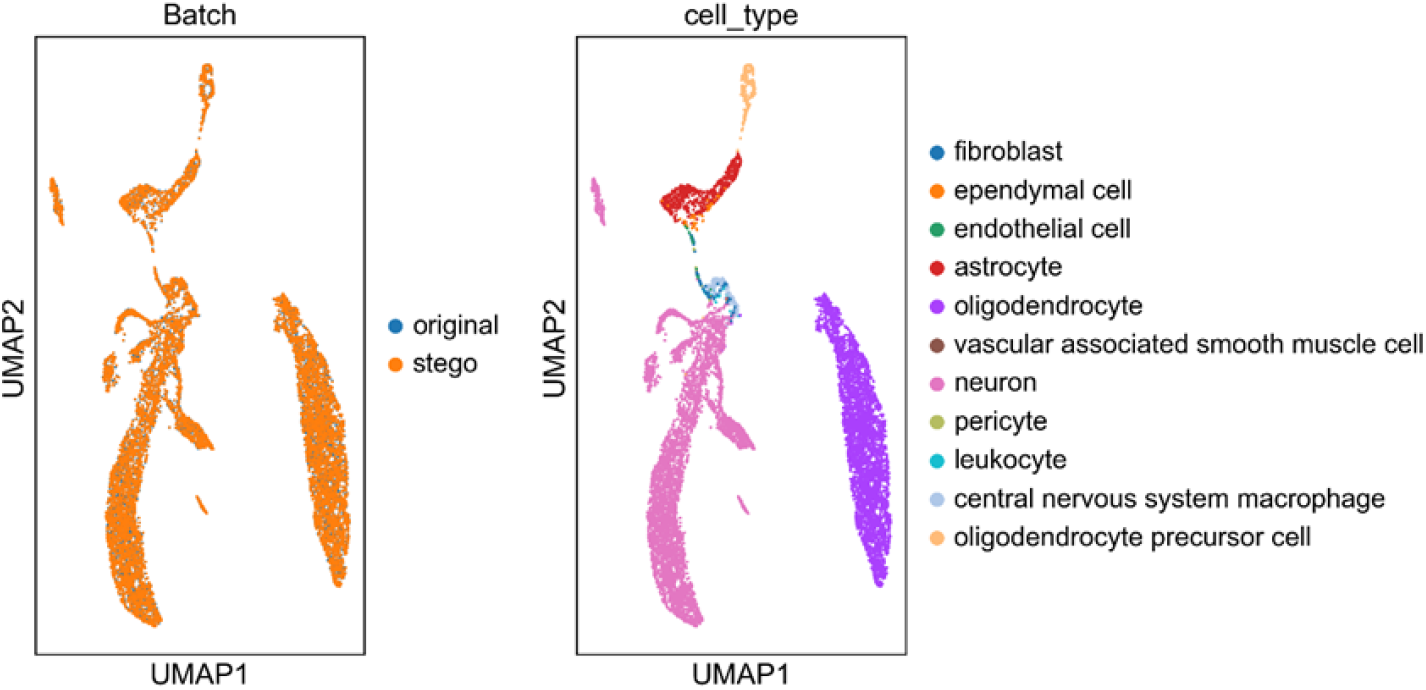
5a3bd24d-2df4-4cf4-b90d-56d457af3ae1_*s*_*n*_3*u*_*mapf igure*

Regarding clustering consistency, unsupervised clustering was performed using KMeans (n_clusters=5) applied to the principal components derived from PCA. The Adjusted Rand Index (ARI) was computed to quantify the agreement between cluster labels obtained from the original and restored (stego) datasets.

In this evaluation, the ARI score reached 1.0, indicating perfect correspondence in cell subpopulation assignments. This result demonstrates that the biological structure of cellular communities, as captured by PCA-based clustering, is fully preserved following data embedding, with no detectable impact on clustering stability or biological interpretability.

### 4.5 Nonlinear dimensionality reduction results

Furthermore, UMAP analysis was employed to evaluate structural preservation under nonlinear dimensionality reduction. The original and stego datasets were concatenated along the cell axis and projected into a shared UMAP embedding space. After PCA reduction to 30 components, a unified neighborhood graph was constructed based on the joint PCA space, and a shared UMAP embedding was generated using consistent parameters (n_comps=30, umap_neighbors=20, mantel_sample_size=1000, etc.), ensuring fair topological comparison between the two datasets.

UMAP visualization revealed that in the plot colored by dataset origin, the original and stego data points were highly intermixed in the 2D embedding space, with no discernible batch separation, indicating that the information embedding process did not introduce significant topological distortion. Meanwhile, when colored by cell type, distinct and biologically meaningful clusters emerged, and the spatial distribution of each cell type was nearly identical between the original and stego data, demonstrating that cell type identification and clustering were unaffected by the embedding procedure. Further quantitative analysis confirmed these observations: the Adjusted Rand Index (ARI) between clustering results of the two datasets reached 1.0, indicating perfect agreement in cluster assignments. Additionally, an approximate Mantel test [26] on pairwise cell distances in the PCA space yielded a correlation coefficient of 1.0, reflecting near-identical global structural similarity. One UMAP figure is shown in 3, more results see [5].

## 5 Conclusion

This paper addresses the challenges of privacy protection in the open sharing of scRNA-seq data by proposing and implementing BioDH, the first reversible data hiding framework designed for gene data. Extensive validation on realworld scRNA-seq datasets demonstrates that BioDH achieves perfect reversibility: both the embedded information and the original expression matrix are reconstructed without any loss. Numerically, the maximum and mean absolute differences between original and stego-datasets are 8.54E-04 and 9.13004E-11, respectively. Biologically, the method preserves global expression patterns with a Pearson correlation of 1.0 on the first two principal components, and UMAP visualizations show nearly indistinguishable low-dimensional embeddings. The approximate Mantel test confirms high structural fidelity (r1.0), and clustering analysis yields an Adjusted Rand Index (ARI) of 1.0, indicating complete consistency in cell subpopulation identification. These results collectively validate that BioDH introduces negligible perturbation across numerical, topological, and biological levels. BioDH balances strong data security with uncompromised scientific utility, fulfilling the stringent requirements for reproducibility and analytical integrity in biomedical research. All code and tools have been made publicly available [5] to promote transparency, community validation, and widespread adoption.

Future efforts will explore higher-capacity methods, enable adaptive compression via lightweight models, and extend BioDH to spatial transcriptomics and multi-omics data—toward a universal, robust, and auditable platform for biological data security.

